# A more-than-human political ecology of Indonesian songbird trade

**DOI:** 10.1101/2025.04.15.648493

**Authors:** Sicily B Fiennes, Novi Hardianto, Silvi D Anaswari, Luthfiyyah Damayani, Jero Haryono, Tom Jackson, Asri A. Dwiyahreni, Christopher Birchall, George Holmes, Christopher Hassall

## Abstract

Since its conception as a discipline, conservation has considered the ‘problem’ of wildlife trade. In focusing on conservation outcomes, we almost wholly omit discussions on the welfare of animals and plants, and the harms they endure. Here, we develop a political ecology approach that incorporates the interconnectedness of people with animals and natural habitats (“more-than-human”) to study the Indonesian bird trade, which is deeply culturally embedded, monetised and speciose. Bringing together marketplace observations, 1-1 interviews with experts, and focus groups with law enforcement, we map out the trade across three levels (actor, inter-actor and market level) to explore flows of birds, interactions, and power dynamics within this economy. We use a method that considers both human and bird perspectives to recognize birds as active participants with their own experiences

Specifically, we acknowledge previously obscured harms experienced by birds like feather plucking, dismemberment, sinus infection, overcrowding, suffocation and death. Different forms of harm occur to birds in different parts of the supply chain and depend on the human actors with whom they are interacting. Loss of freedom occurs at harvest and physical/physiological harm during transit and at the point of trade. However, harms are lower for highly sought-after species, though they are difficult to source and are well cared for by affluent collectors, but harms are higher when demand is high, and supply-side factors lead to broad harvesting with lower consideration of welfare. Our findings also indicate that men of different classes engage with birds for various reasons, such as for socialisation, investment and connecting with Javan traditions. We use this interdisciplinary approach to highlight the harms birds experience during the wildlife trade, relating to the Five Domains welfare model. Critical to understanding the harms endured are issues surrounding class, gender and culture in Indonesia, and other IWT contexts.

## Introduction

Global capitalism and international trade have formed flows of animals, plants and fungi (Hodgetts and Lorimer 2015), commonly referred to as legal and illegal wildlife trade (IWT) (Phelps et al. 2016). Some claim wildlife trade is an ‘extinction market’, where species are trafficked to extinction (Felbab-Brown 2017), though the extent to which wildlife trade drives extinctions is unclear, especially since many species face habitat fragmentation and climate change in synergy (Hinsley et al. 2023). As such wildlife trade is predominantly seen as the concern of conservation by policymakers, conservation organizations, and regulatory bodies. Wildlife trade has been justified on the basis that it can be legal, safe, sustainable and traceable. Others argue against wildlife trade, based on its public health risks, such as zoonosis (Yang et al. 2020). IWT, a narrow set of activities in breach of national-level regulations (Duffy 2022, p.32), usually connotes trade in rare species or at unsustainable levels. However, IWT is not always detrimental to wild species and recent evidence suggests that legal trade can be just as harmful and unsustainable (Marshall et al. 2025).

Discussions about wildlife trade within conservation omit the materiality and experience of the individual entity that is trafficked, and how their life is transformed as they enter the wildlife trade circuit (Collard 2014). This omission is partly due to conservation’s normative commitments to nonhuman animal populations and their habitats (Campbell 2012), and its imagination of animals as passive taxonomic objects. Affect for individual animals is often seen as the remit of animal welfare advocates and animal rights activists, though both conservation and animal rights movements have been critiqued for promoting unjust and uneven politics in the name of non-human lives (Oommen et al. 2019). Rather than recognising the trafficked animals as agents, both conservation and wildlife trade see animals as objects, properties and commodities rather than subjects, individuals and sentient beings, whether they are objects to be conserved or sold (Wyatt et al. 2022). Viewing wildlife as objects has three key limitations for a fuller understanding of wildlife trade: it ignores the social and cultural ideas that shape trades, it neglects how the material properties of the animals influence how trade happens, and it marginalises the welfare implications of trades.

The physical properties and behaviours of animals shape how they experience trade and harm but also shape markets in turn. For example, Dickinson (2022) shows how the fleshy material properties of caviar enable its laundering. Its chemical-isotope profile (over time the flesh of illegally laundered sturgeon can develop similar characteristics to that of captive-bred sturgeon) and composite form (caviar eggs are small and when thousands of legal and illegal eggs are mixed in tins it is impossible to verify their origin) contribute to its existence as a grey trade (Dickinson 2022).

The exploitation of less charismatic wildlife for food, as pets, and for recreation is a major driver of wildlife harm (Hutchinson 2023). One complex case of wildlife harm is the caged bird trade in Indonesia, the global hotspot for bird trade. A segment of the market caters to Javan traditions that dictate men need various possessions to constitute a ‘good life’, one of which is a kukila (cockfight bird or songbird) (Yulindrasari 2017), though the practice is also influenced by royal nobility who used to display elite species outside their palaces (Pradita and Wardhana 2021). These regal practices formed the foundations for another subculture where birds represent status symbols.

Emerging out of these older practices and formally materializing in the 1970s (Mirin and Klinck 2021), singing competitions involve the training of various species of songbirds to sing and outcompete other birds. The commercialisation of singing competitions has meant that birds now provide important real and imagined animal capital (Hodgetts and Lorimer 2015), not just by elevating social status or engaging with Javanese traditions but also through instrumental and relational value (i.e. selling rare and expensive birds, selling competition winners and earnings from singing competitions). Birds’ capital is intrinsically tied to their materiality; they are easy to transport and hide because of their small size (Maulany et al. 2021), and Indonesia’s high avian diversity ensures that one species is fungible for another (Fiennes et al. 2024).

Pons-Hernandez (2024) argues that ‘it is easier to exploit (trade, traffic, or eat) nonhuman animals if we do not see their suffering’. A harms-based and victim-centred approach to wildlife trade (Wyatt et al. 2022; Hutchinson 2023) can help us understand the scale and nature of animal suffering but also understand what these harms reveal about the underlying structural drivers of IWT (Duffy 2022). Multispecies ethnography (MSE) provides a tool by which to study the human and non-human actors in IWT while recognising animals as subjects, individuals and sentient beings (Wyatt et al. 2022). Multispecies ethnography has focused on the many individuals of one species; see Tomlinson (2020) on horses or mice, or Tsing (2015) on the matsutake mushroom. MSEs of live wild animals in the wildlife trade, however, are scarce.

Here we build on Oyanedel et al.’s (2024) seminal framework to understand macroeconomic markets for wildlife as a complex system of actors, market dynamics, and overall market operation, by considering the interconnectedness of human life with non-human entities in wildlife trades. This allows us to incorporate birds into the trade system as nonconsenting actors. By following their journeys in a ‘follow-the-thing’ sense (Dickinson 2022), we can gain insights into how birds’ physical properties and behaviours influence their experiences. We treat the wildlife trade as a cultural phenomenon, influenced by social differences like gender (Margulies et al. 2023), class and race (Margulies et al. 2019) that shape harms committed to traded animals. Here, we explore the harms done to individual humans and non-humans within the wildlife trade and how this is shaped by human culture and the material properties of animals involved.

## Methods

We draw from five months of fieldwork (January to June 2023) across Java (Jakarta, Bandung, Yogyakarta, and Surabaya) and West Kalimantan (Pontianak) in Indonesia. This fieldwork involved three methods to conduct a more-than-human political ecology of the Indonesian songbird trade: short multispecies ethnography at bird marketplaces, focus groups with conservation law enforcement officers, and interviews with conservation practitioners and veterinarians.

### Visiting wildlife marketplaces

We visited nine wildlife marketplaces across each of the five research locations, with each marketplace visited at least twice, except for one in Jakarta, which was visited once. There are inherent tensions between creating ethnographic accounts that are empirically thick and the consequences of making people and places less anonymous (Lyon and Back 2012). To remedy this tension, we do not refer to marketplaces by name, nor do we publish images where the marketplace could be identified.

We take inspiration from the traditions of sensory ethnography and short ethnography to conduct a short multispecies ethnography (MSE). Short-term ethnographies are a focused and time-efficient approach to classical ethnographic research using immersive and participatory methods to provide a deep understanding of places (Pink and Morgan 2013). We use sensory ethnography methods to get a better understanding of the experience of nonhuman animals (Tomlinson 2020). We adopt a form of MSE that treats nonhuman animals as intersubjective, individual actors who experience pain and emotions (Tomlinson 2020). Building on ethnographic methods of ‘hanging out’, we employed ‘hanging around the market’ to absorb the sensory and architectural environment of the marketplace. By visually documenting trade, which can enhance understanding of social practices and experiences during a condensed research period (Pink and Morgan 2013), we provide evidence of harms (Pons-Hernandez 2024). When we refer to specific species, we follow the taxonomy listed in the BirdLife DataZone.

### Focus groups, interviews and human consent

In each city, we held focus groups (n = 5) with Indonesian conservation law enforcement agencies (57 participants from these agencies took part, with 8 to 13 participants per session) to discuss their roles and challenges in enforcing songbird IWT across the supply chain. We collaborated with four agencies: BKSDA (nature conservation), BKP (animal quarantine), GAKKUM (wildlife law enforcement), and POLRES (regional police). These agencies are referred to with the abbreviations BKSDA, KAR (Karantina = Quarantine), GAKK and POL. We conducted sixteen semi-structured interviews (eight in person, eight online) with experts from conservation organizations (represented with NGO in participant codes) and universities (using ACA for academic in participant codes). Interviewee codes are followed by an L for local or an I for international. These interviews focused on bird trade as a system and interviewees were asked to provide feedback on a preliminary supply chain diagram. All participants were compensated at a standard hourly rate and gave their free, prior, and informed consent (FPIC).

We analyzed qualitative data from marketplace observations, focus group discussions, and interviews using NVivo 11. Focus groups were conducted in Indonesian, recorded, transcribed, and translated (see Supplementary Information Guide 1). Interviews were conducted either in English or Indonesian, transcribed and translated into English if needed (see Supplementary Information Guide 2).

### Vignettes and welfare assessment

Birds cannot give their consent to being photographed and surveilled. By being a spectator we risk reinforcing anthropocentric hierarchies (Hooper et al. 2022). During interviews with veterinarians, we use vignettes to critique our own anthropomorphism and provide a reflective exploration of the implications of projecting human qualities onto non-human entities, and the difficulty and limits of accessing animal subjects’ experiential worlds (Hodgetts and Lorimer 2015).

We discuss a series of images and videos that depict avian suffering in marketplaces with veterinarians and present them here as vignettes. Vignettes are useful since they describe information in context (Rizzolo 2021), and can be used to clarify judgements. We use these as a source of expert opinion and categorise the resultant harms according to the Five Domains model (Mellor et al. 2020). This model acknowledges that animals have psychological as well as physiological needs. Bartels et al. (2022) used an older version, the Five Freedoms, to assess the welfare of captive-bred birds at marketplaces in Germany (both models are compared in SI Table 1). We consider the Five Domains in the following ways:

1. Nutrition: differences in diet in the wild vs. in trade
2. Environment: contrasts between the natural habitat with the confined spaces
3. Health: any health impacts, potential injuries, and lack of medical care
4. Behaviour: examine stress behaviors or inability to exhibit natural behaviors in trade settings
5. Mental State: explore how changes in the first four domains affect the bird’s mental well-being.

**Table 1.** Examples of violations of the Five Domains towards birds in trade, including from vignettes and other marketplace observations.

| Domain | Example |
| --- | --- |
| Environment | Overcrowding (SI Figure 1A) |
|  | Birds covered in/ in close proximity in excrement (SI Figure 1C) |
|  | Cages left in direct sunlight (Vignette 3) |
| Health | Broken limbs and damaged feathers (Vignette 1) can result from birdlime use |
|  | Infections (sinusitis) from dust in the marketplace sinusitis (Vignettes 3 and 4). |
| Behavior | Being active, trying to fly away or escape (Vignettes 3 and 5) |
|  | Plucking feathers on their back from stress (Vignette 2) |

### Assessing the Indonesian macroeconomic market driving unsustainable bird use

We structure the results of our more-than-human political ecology through the three stages of Oyanedel et al.’s (2024) framework. The multispecies ethnography and vignettes contribute to our understanding of birds, welfare issues and how birds are moved in supply chains. The interviews, together with the focus groups, allow us to assess power dynamics within humans and between humans and animals. In combination, these methods allow us to understand the harm to birds and people within this system, based on social framings like power. We integrate birds across three analytical levels (actors, market dynamics, and overall market operation) and add an economic scale: microeconomic, mesoeconomic, and macroeconomic.

#### 1. Characterization of songbird trade actors (activities at the microeconomic scale)

We identify human actors in the caged bird trade, including harvesters, intermediaries, vendors, and consumers, considering cultural dynamics. We corroborate actor roles with existing literature, noting the multitude of roles in IWT market chains (Phelps et al. 2016). Birds are established as non-human actors, detailing their transition from wild to commodity lives using the Five Domains model.

#### 2. Examination of interactions (transactions at the mesoeconomic scale)

We focus on the relationships and interactions between market actors (human and non-human), the structure of supply chains, and the methods used to harvest and transport free-living wild birds to caged environments.

#### 3. Assessment of market dynamics (market level, macroeconomic scale)

Lastly, we consider broader market dynamics such as supply-demand interactions, legal, grey and illegal entanglements and broader human-bird relations in Indonesia.

We visually summarise existing information on 1. how human actors are arranged and their relations, 2. how sourcing pressure has been displaced from Java, the core hotspot of trade, to Sumatra and West Kalimantan and 3. the legal, grey and illegal entanglements that bring birds to marketplaces.

## Ethics

We received ethics approval from the University of Leeds (AREA FREC 2023-0419-521) and fieldwork was conducted under a research permit (311/SIP/IV/FR/11/2022) from Indonesia’s National Research and Innovation Agency.

## Results

The songbird trade network is made up of two parallel supply chains (Figure 1A-C), aligning with those proposed by Jepson et al. (2011)’:*1. local people and 2. syndicates taking birds from forests on a large scale, involving established business elites’* (ACL1). Longer supply chains may be utilised more for high-volume and low-worth species, catering to entry-level hobbyists and breeders, and the species composition is more random, supported by opportunistic harvesters. Shorter supply chains will be utilised for higher-end species (rare and/or expensive) or for orders placed by consumers (Figure 2A). There are some interactions between the supply chains, for instance, breeders are now also turning their hand to expensive and rare species.

**Figure 1.**
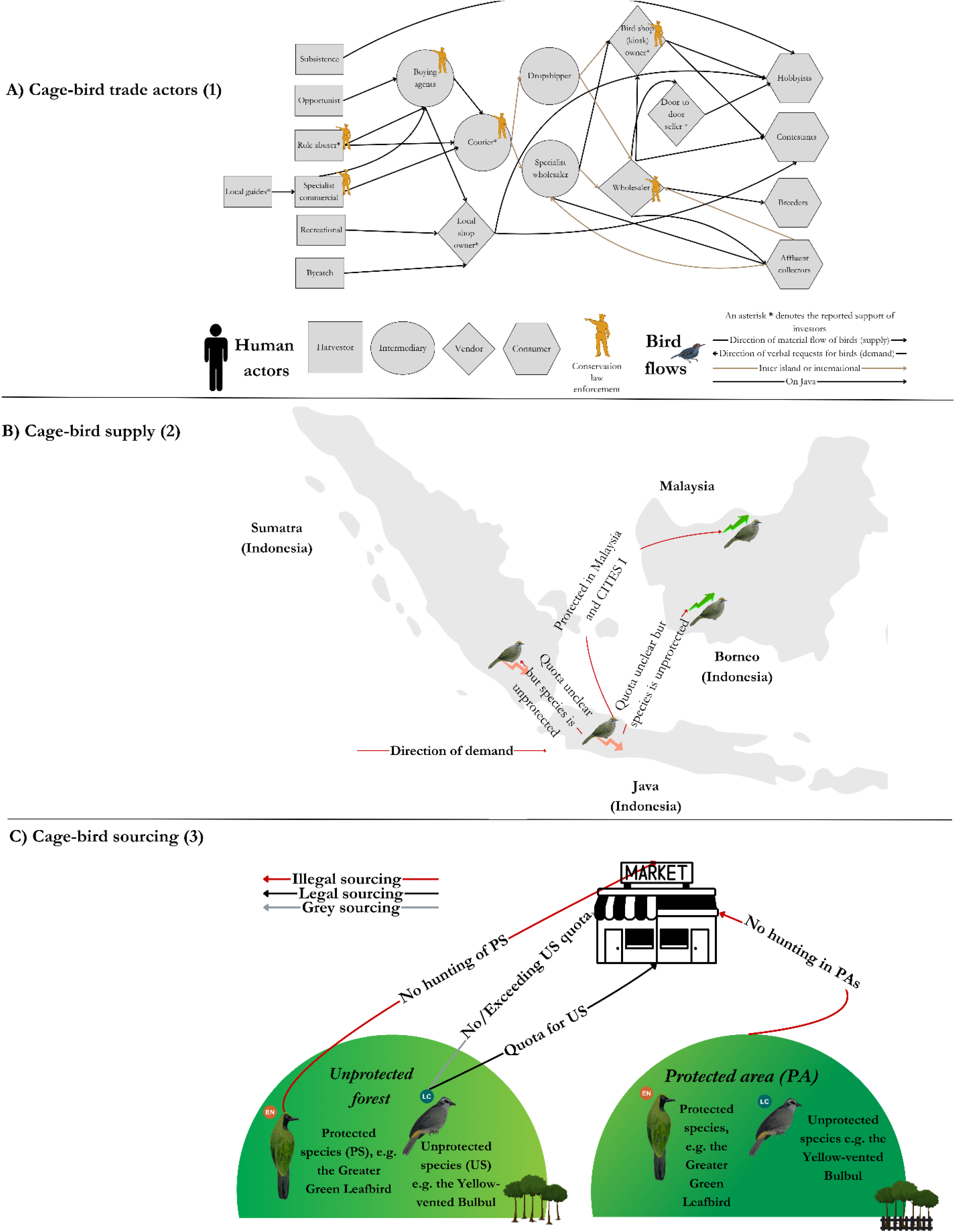
**A):** Actor map depicting the flow of birds through four stages: harvesters, intermediaries, vendors, and consumers. Quotes in SI Table 3 support the roles of each actor. **B)** Local extinction on Java and Sumatra drives inter-island (Kalimantan) and international trade (Malaysia), represented by the Straw-headed Bulbul, *Pycnonotus zeylanicus*, with range data based on Chiok *et al*. (2020). Birds drawn by Ishaan Patil. **C)** Pathways (legal, grey, illegal) for supplying birds from various sources (unprotected/protected areas) to markets, illustrating the structure and dynamics of the supply chain.

**Figure 2.**
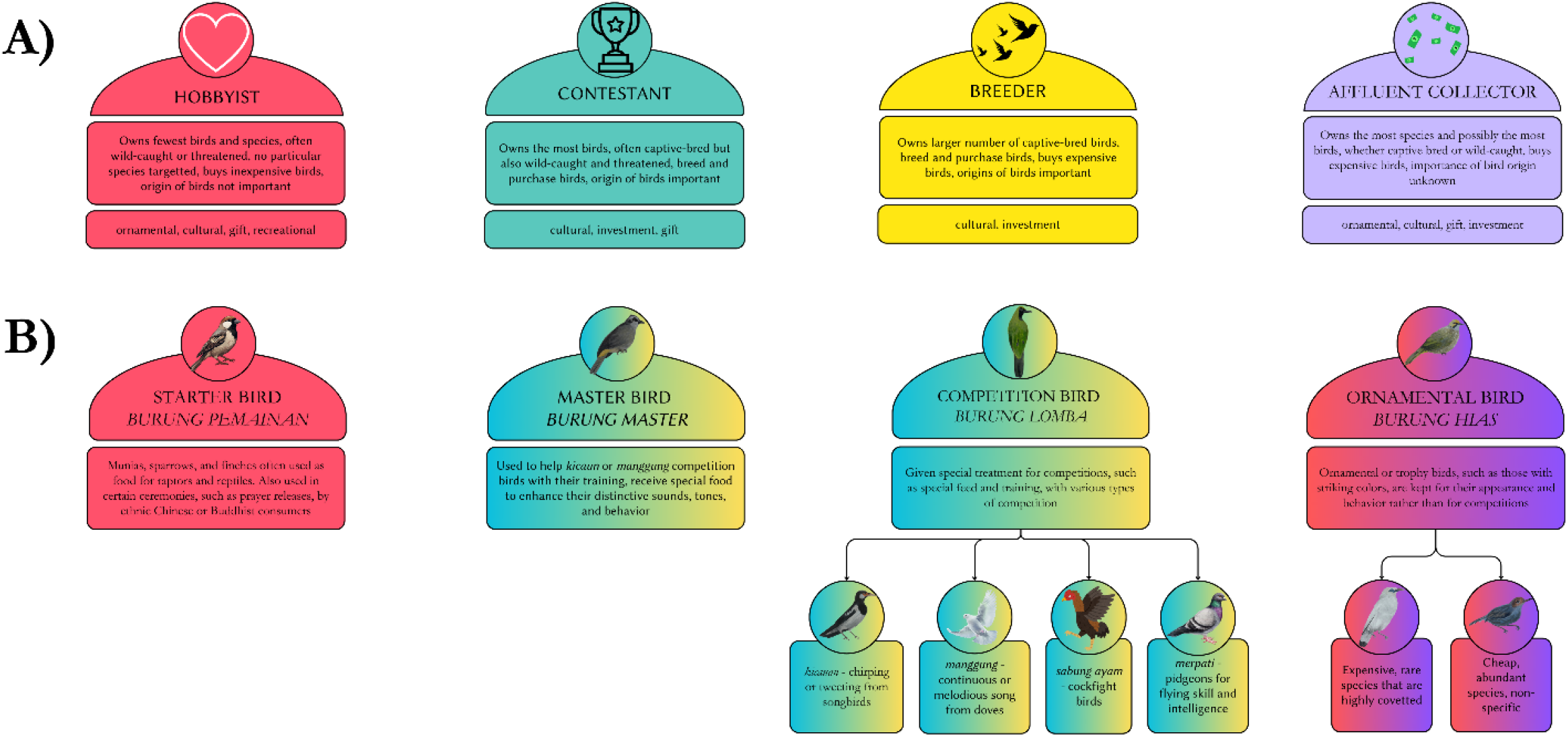
**A)** User groups in the Indonesian bird trade, including 1. Marshall et al.’s (2020) groups: hobbyist, contestant, and breeder, and our new category; affluent collector, 2. consumer uses from Phelps et al. (2016). **B)** Bird groups: house birds, master birds, competition birds and ornamental birds, overlap between human user groups with birds can be seen in the hybrid colours. Inspiration for some of the bird groups is taken from broader categories from Jepson et al. (2011) and Basuni and Setiyani (1989).

One interviewee estimated that over 80% of songbird trade in Indonesia is illegal (i.e. protected species, trapped in protected areas or exceeding quotas (Figure 1C) (NGOL7)). As we will go on to demonstrate, trade is decentralised; partly opportunistic, and partly organised (Figure 1A-C). Therefore the Indonesian wild bird trade is ‘crime that is organised’ rather than ‘organised crime’ (distinguished by Pires et al. (2016), when referring to wildlife crimes).

Next, we present novel insights into the unsustainability of the Indonesian bird trade, focusing on actors, inter-actor dynamics, and the broader bird economy, particularly around aspects that help us understand the harm committed to birds.

### 1. Characterization of bird trade actors

We detail the emergence of an undescribed user group, the affluent collector (Figure 1A, 2A). The consumer roles suggested by Phelps et al. (2016) also relate to the aesthetic and cultural values placed on songbirds by consumers because they are beautiful or seen as trophies (ornamental birds) or to fulfil deeply entrenched Javan cultural values (*manggung* or *merpati* competition birds). Other bird groups exist because of their market-based instrumental value (Hutchinson 2023), heavily influenced by consumer cultures (Thomsen et al. 2024), noticeably *kicauan* and regal practices (expensive ornamental birds) (Figure 2B).

‘*Apart from traditional* [sabung ayam, manggung or merpati] *or well-known good birds [high-end ornamental species like the Straw-headed Bulbul], trends are likely dictated by traders and availability [in the wild]’* (NGOI1). Other work (e.g. Marshall et al. (2020); Indraswari et al. (2024)) has shown that the contestant and breeder community demand a relatively fixed suite of *kicauan* species that are eligible for singing competitions.

A single consumer may own various bird profiles, from low-value ornamental birds for hobbyists to master and competition birds for contestants and high-value birds for affluent collectors. Bird profiles can overlap, as one person’s master bird may be another’s ornamental bird (Figure 2B).

#### Vignettes with veterinary experts

Attending to the experiences of nonhuman animals is fundamentally a challenging task as ‘*wild animals have a masking phenomenon… They hide sickness to avoid predation’* (NGOL3). These vignettes highlight common misunderstandings around how animal harms are understood, particularly in the health, behaviour and mental states domains. In almost every instance, the initial interpretation of the authors was incorrect. For instance, the Sunda Scops-owl, *Otus lempiji* (Vignette 4) most likely has a sinus infection, though it initially looked like a trauma wound.

**Figure V1.**
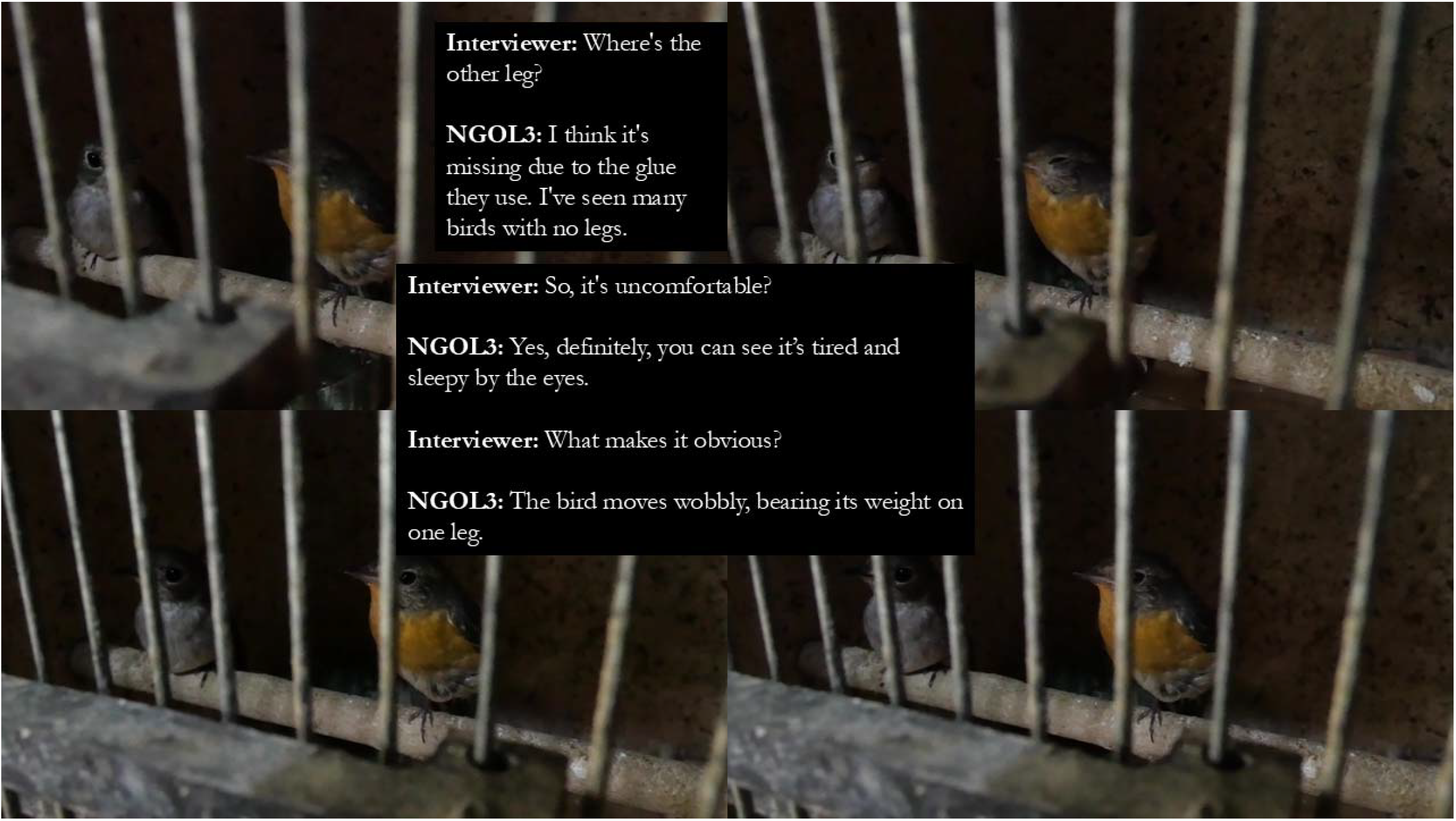
Asian Brown Flycatcher, *Muscicapa latirostris* (left) and a Mugimaki Flycatcher, *Ficedula mugimaki* (right).

**Figure V2.**
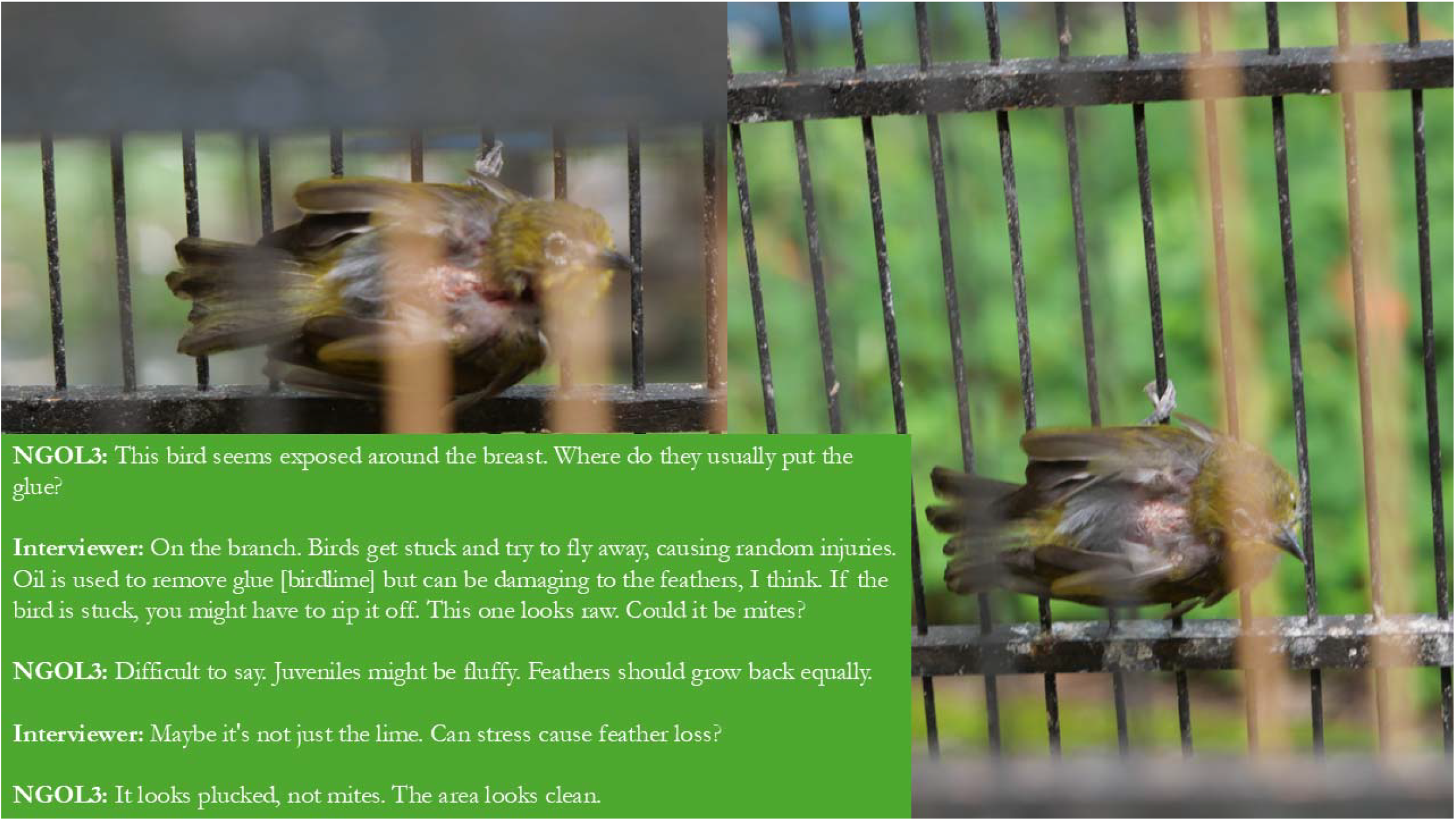
a Hume’s White-eye, *Zosterops auriventer*.

**Figure V3.**
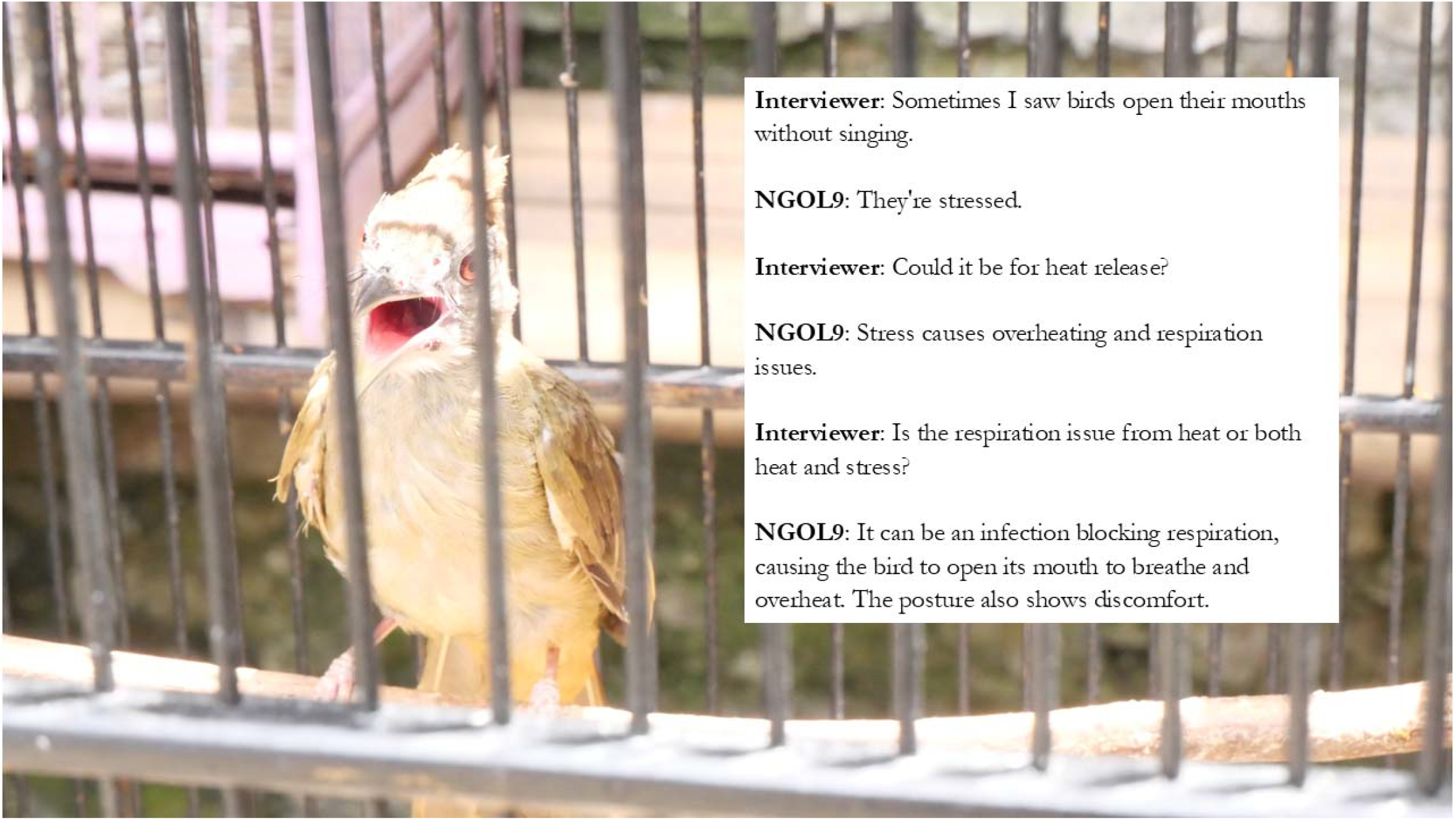
an Ochraceous Bulbul, *Alophoixus ochraceus*

**Figure V4.**
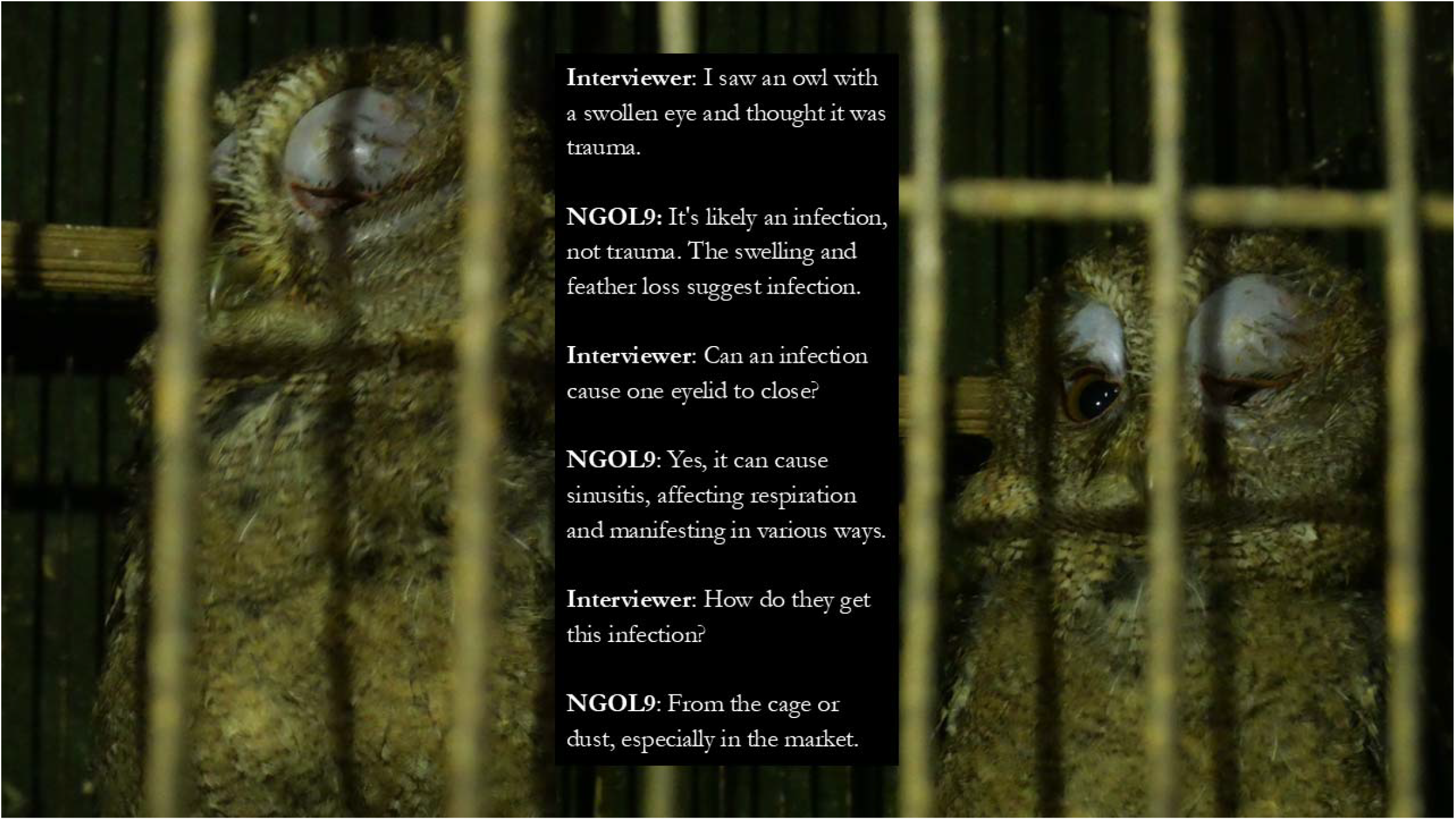
a Sunda Scops-owl, *Otus lempiji*

**Figure V5.**
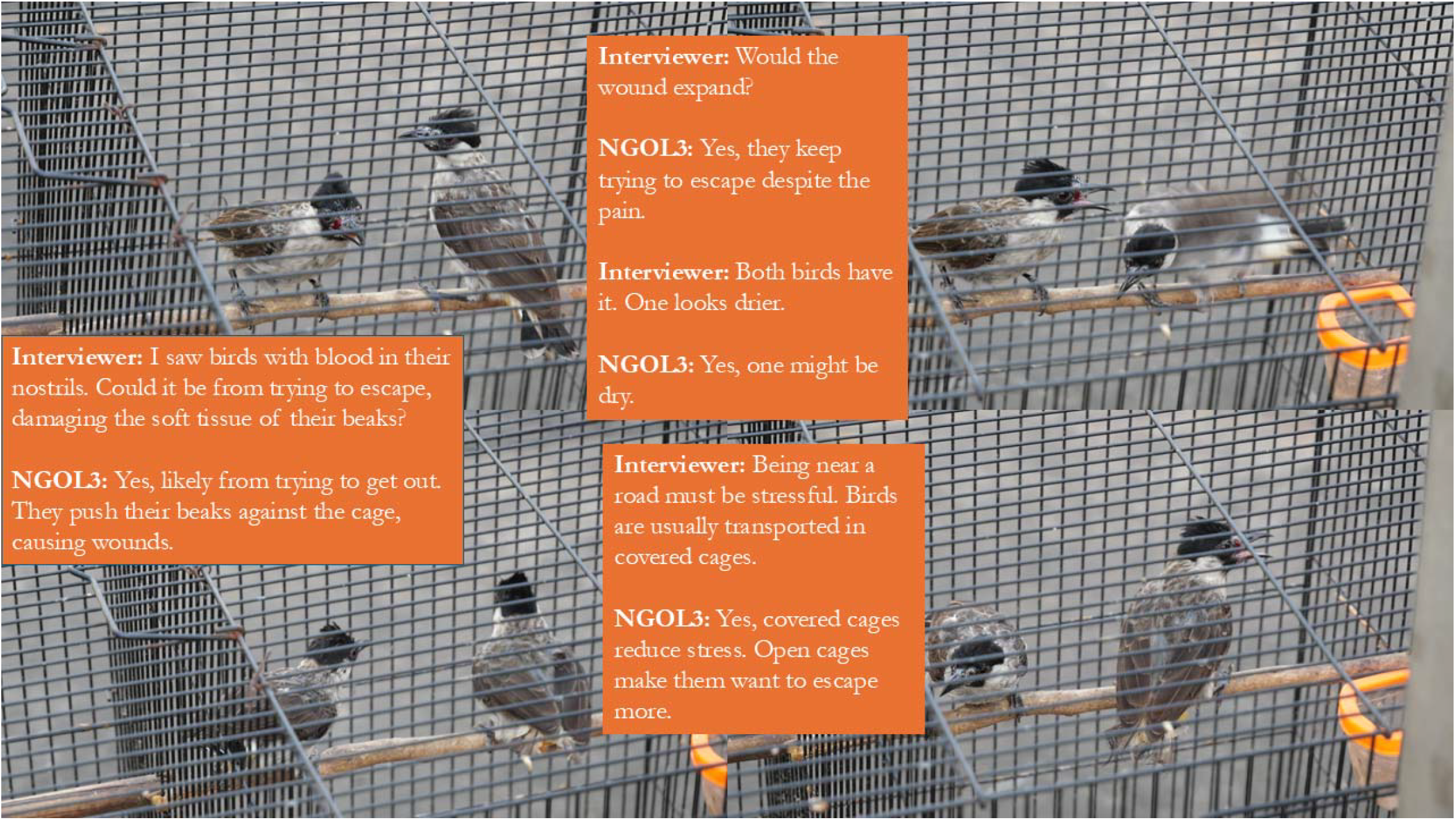
a pair of Sooty-headed Bulbuls, *Pycnonotus aurigaster*.

#### Five Domains: What is done to birds?

Even after taking a more critically anthropomorphic approach, it is evident that the fleshy properties of birds are changed, altered and removed in the marketplace (Table 1). In SI Table 2, we present the same information, along with additional violations noted in the literature and hypothesized differences between the wild and captive lives of birds.

### 2. Inter-actor analysis

#### 2.1. Cooperation, ordering and financing

Demand from consumers does not always drive supply. If a vendor receives a request in-person or an order, they may cooperate with other vendors: *‘If his shop sells out of White-rumped Shamas and I have some, I can sell them to him’* (NGOL8). However, interviewees also suggested that more influential actors like wholesalers finance smaller-scale vendors like kiosk owners or door-to-door sellers, as well as external investors (denoted with an * in Figure 1A), so some vendors may have little control over their supply.

Given there has been a massive and difficult-to-quantify drain on Java’s wild bird populations, most trade may now be inter-island (Figure 1C), made possible through the use of passenger ships (ACL1, GAKK_6, GAKK_9). ‘*[For orders] they use private vehicles’* (NGOL8, GAKK_8, GAKK_9), ‘*to reduce deaths. Deaths mean losses’* (NGOL6).

For the supply of expensive ornamental birds like *‘parrots… [they use] navy vessels or special aeroplanes’* (GAKK_8) and the modus operandi can be tailored to evade detection by authorities, such as exchanging birds at sea (GAKK_8, BKSDA_17) or going through non-public ports (GAKK_9).

The quality and condition of birds in the marketplace depends on how far away the bird is sourced; for the main inter-island trade routes (i.e. South Sumatra to northern Java and West Kalimantan to East Java, via Madura (GAKK_8, GAKK_9)), the journey could be half a day or several days if birds come from further afield (e.g. Papua (SI Figure 2)).

#### 2.2. Methods for harvesting and moving birds

Bird harvesting draws on knowledge of avian culture to adapt methods to bird behaviour (Lappe-Osthege and Duffy 2024). In the Indonesian songbird trade, this can be seen in the use of playback on speakers to lure birds using decoy birds (Basuni and Setiyani 1989), capitalising on male-male competition or relying on intricate methods derived from other artisanal practices like fishing (Figure 3, SI Table 4). How birds are trapped is also linked to the resources available to the trapper: ‘*bird lime is traditional and cheap’*, whereas ‘*mist nets are quite expensive, costing around USD* 50’ (NGOI2). Birdlime has higher mortality rates, and birds sell for less (NGOL7), and as we have shown (Vignette 1), it can lead to dismemberment.

**Figure 3.**
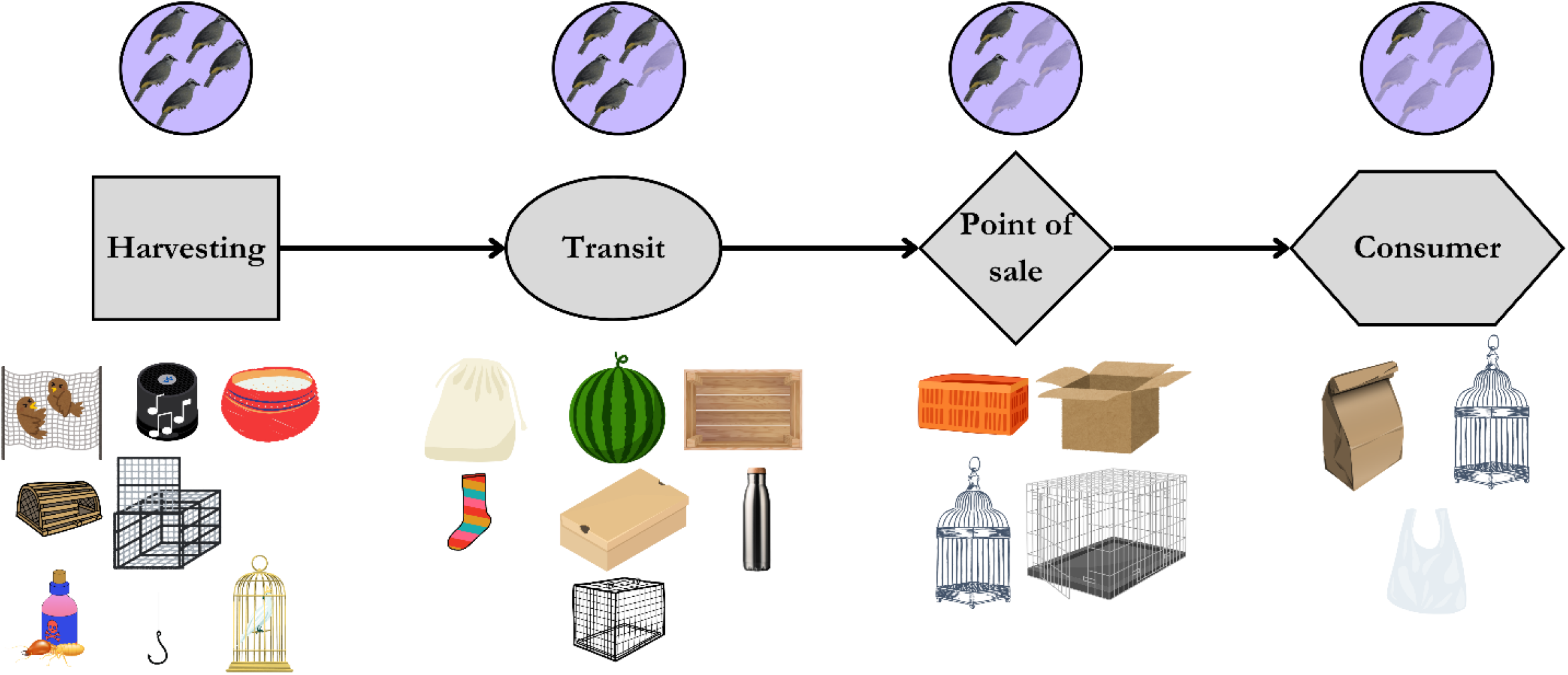
Summary of methods for harvesting and transporting birds for the live pet trade in Indonesia. Peng and Broom report a 90% mortality rate for small birds captured for pets. With 1 million birds trapped annually, 233,333 survive to be sold, indicating a 77% mortality rate. This hypothesis is based on incomplete data for small-bodied, sensitive species. Shepherd et al. (2004) found a 50% mortality rate for munias in the first 24 hours after capture. Confiscation data shows a 33% mortality rate during transit (Ramani and Clark-Shen 2019). Marketplace conditions further reduce survival rates. Approximately 1 in 4.35 birds harvested will survive. Starting with 5 birds, 1 will survive to reach the consumer.

### 3. Assessment of market dynamics

#### Human-bird relations in marketplaces

Though mortality rates may be high across the supply chain (Figure 3), these, along with welfare standards, are dependent on the bird group. Extensive welfare violations and higher mortality rates are seen for small-bodied species like the munias (Table 1, SI Figure 1B/D), from the casual bird and cheap ornamental bird groups (Figure 2B).

There are various techniques used to advertise birds to consumers, usually for low-cost species. Some birds were traded in cages branded with international football team logos; one trader named a bird the ‘Arsenal bird’ (SI Figure 4). Other casual birds are spray-painted to be marketable to children (SI Figure 1 E)). These techniques may help to sell ‘*new and unknown species by making them more appealing and exotic’* (NGOL4), but for poorer vendors, there is a trade-off between keeping inexpensive birds alive versus letting them die.

Shops specialising in expensive ornamental species (e.g. rare native species like the Straw-headed Bulbul, *Pycnonotus zeylanicus*, or prized songbirds from China like the Black-throated Laughingthrush, *Garrulax chinensis* and Hwameis, *Garrulax canorus*) and popular *kicauan* competition birds (White-rumped Shama, *Copsychus malabaricus*, Oriental Magpie-Robin, *Copsychus saularis*, and the Orange-headed Thrush, *Geokichla citrina)* were much cleaner. Birds from these groups were sold in larger, excrement-free cages with varied food, and customers were asked to remove their shoes before entering the shop. ‘*High-priced, legally risky, or difficult-to-care-for birds tend to stay longer’* (NGOL6), but that is also because the vendor can afford to look after those birds.

Poorer consumers have fewer resources available for multispecies care (Bubandt 2024) and are unable to provide their birds with the best husbandry. Despite financial limitations, birdkeepers create home remedies based on community knowledge, often avoiding veterinary visits (NGOL4) and *‘since COVID, there has been a rise in people building larger aviaries, allowing birds to spread their wings and fly, which is a significant improvement’*

(NGOI1). Further, vendors can still sell sick birds, and the price may be related to how well the species survive. This provides additional clarity on the “cut flower” phenomenon noted amongst low-level hobbyist consumers (e.g. bird mortality is highest amongst hobbyists (Marshall et al. 2020). It might not be that they have the poorest level of husbandry but that they are buying cheap birds that are either sick already or sensitive to the conditions along the supply chain.

## Discussion

We reveal that the Indonesian songbird trade is highly complex, informal and comprised of several overlapping pathways of supply. We detail harms to birds including infection, dismemberment and stress behaviours. However, these harms are mediated by the social differences of human actors (we explore those of vendors and consumers here), as well as the bird group. Multiple perspectives and value systems are inherent to wicked problems and IWT. It is likely extremely difficult to change how wildlife is valued and it is evident that songbird trade and associated harms to birds would vary if these cultural aspects were different. We contextualise our findings within the literature on other wildlife trade and small-scale economies and reflect on the questions of power (class relations, labour precarity, etc) that explicitly dictate the harms birds face.

### Cooperation, investment and power

It has been suggested that most wild bird harvest in developing countries is a “cottage industry” (i.e. production in the home) (European Food Safety Authority 2006). Here, we show that wild bird harvest is not a cottage industry but a series of micro-enterprises that form artisanal clusters, catering to low-income consumers, where the role of intermediaries and vendors is dominant (Tambunan 2005). The growth of the Indonesian songbird trade is constrained by the reliance of poorer actors on powerful vendors such as traders, through informal and unequal financial relations.

The poorest human actors are likely harvesters. Bird harvesters (for the food market) needed to borrow from local wholesalers to support hunting activities in Java and Sumatra (McCarthy and Noor 1996), which ties them to local markets. There is precarity from economic instability but also from physical risks to actors who provide labour at the site of production for birds (i.e. the forest). Diminishing bird populations drive harvesters deeper into forest interiors, sometimes involving walking more than three hours on foot to camp for several days and carrying birds back (Lucas 2011). The increase in illegal logging and agricultural encroachment in Sumatra and Kalimantan created new sources for wild-caught birds (Jepson et al. 2011) but also puts harvesters at risk of encountering snakes, tigers and bears (Silalahi 2020). This precarity is also seen in Romania’s timber trade (Iordăchescu and Vasile 2023) where there is a class of precarious forest workers and petty woodcutters.

Turnover is not always high for rare species, and vendors are absorbing the costs of keeping animals alive as well as competing with online sources like dropshippers (someone who takes customer orders and forwards them to a supplier, typically through an online store) (Figure 1A). In surveys with bird traders across West Kalimantan, Miller et al. (2019) found that for 40% of bird shop owners, it was not their sole income source, nor was it particularly lucrative. Evers and Mehmet (1994) surveyed small-scale agricultural traders in Java, funding that nearly half borrowed money for business, mainly from informal sources like moneylenders. Wholesalers (traders) and dropshippers are likely to be financially independent and able to offer ordering options to wealthier consumers (Figure 2A). Black-winged Myna, *Acridotheres melanopterus*, that were stolen from a conservation breeding centre in West Java were all sold at bird markets in Jakarta the next day, which indicated that the birds were pre-ordered (Sozer and Tritto 2014).

### Songbird human actors: understanding the intersections of gender and class

The structure of bird trade reflects the broader informality of Indonesia’s economy (Hao and Freischlad 2022), though as Thomsen et al. (2024) notes, birds and other wildlife are reduced to objects in part by the prevailing neocolonial capitalist framework, where developing nations like Indonesia are left little option but to exploit their own natural resources to partake in global markets. We advance understanding of the gendered consumption of birds (Miller et al. 2019; Bubandt 2024), showing that the class of the consumer relates to how birds are treated (i.e. their welfare), as well as how consumers interact with other trade actors.

Marketplaces are more than just transactional sites; they are cultural environments used as social spaces by trade actors, particularly hobbyists, contestants, door-to-door sellers and bird kiosk owners. Alcano (2016) observed the importance of street-side male socialisation and aggregation in low-income neighbourhoods in Surabaya, East Java, around activities like pigeon racing and gambling.

Bubandt (2024) suggests that songbird keeping evokes nostalgia for village life, fostering male companionship. However, we also highlight the coming together of modern hobbies like football fandom (SI Figure 4) with bird trade, reflecting contemporary social bonds between men. There are parallels between birdkeeping (kicau mania) and Indonesian football culture (bola mania)(Maulida 2024) that both involve ritualised, male-dominated activities, showing the global and modern aspects of these traditional social spaces.

Gender (specifically *cis*-men) plays a significant role in other trades, like cacti and succulent trade, and aphrodisiac trades (e.g., rhino horn or sea turtle eggs). In cacti and succulent IWT, Margulies et al. (2023) hypothesise that men may be driven by heteromasculine notions of being daring. Conversely, Bubandt (2024) suggested that men see songbird competitions as safer than illegal activities like cockfights, even if the competition birds are illegally sourced. Marshall et al. (2021) reported a minimal fear of prosecution relating to songbird keeping amongst Javan consumers. Our results suggest that certain groups of birds are important as a medium of socialisation, something Sánchez-Mercado et al. (2020) noted in the international red siskin bird trade, where breeders, mainly middle-aged men, sought peer recognition most of all.

Our findings also indicate that men of different classes engage with birds for other reasons, such as for investment and connecting with Javan traditions. Poorer contestants (Figure 2A) may view *kicauan* competition or ornamental birds (Figure 2B) as an investment, as a means of class mobility (Lowen 2016). The urban middle class may aspire to be Javanese princes, and expensive ornamental birds once favoured by princes form part of that imaginary (Bubandt 2024). Affluent collectors can reinforce their social status by amassing diverse menageries of species.

### Multispecies structural violence of the cage-bird trade

Relationships between human actors and the value of the bird aside, there is still structural violence embedded in the cage-bird trade. We document several methods of trapping and moving birds (Figure 3) of varying degrees of cruelty, ranging from the use of fishhooks, termites impregnated with poison and watermelons for transporting birds. These methods bear heavy similarity to fishing and hunting birds for meat. In Indonesia, Jepson et al. (2001) noted that Tanimbar Corellas, *Cacatua goffiniana*, were snared with nooses made from nylon fishing lines. Birdlime is used in the European songbird meat trade (Lappe-Osthege and Duffy 2024) and elsewhere in Southeast Asia, such as Myanmar (Platt et al. 2012).

Birds’ behavior influences their trade experiences. Prey species, like songbirds, hide their pain (Doneley 2011). This is a critical factor in understanding why their exploitation is normalised, with minimal recognition of individual harms, which Hutchinson (2023) also observed in the European songbird meat trade. In the case of slaughterhouses, Hodgetts and Lorimer (2020) argue that the architecture and carcerality of the cage disrupt an animal’s ability to express their subjectivities and thus human efforts to understand them.

Though birds’ properties limit our ability to ‘think like a bird’ in complex and noisy marketplaces, a visual inspection is often all that is possible to document the harm towards birds. This approach was also fruitful for Pons-Hernandez (2024), who, in a series of photographs, revealed new and unexplored harms in another high-volume animal trade, the eel trade. These included drawing attention to the risks of eels suffocating through means of transport like plastic bags and providing evidence of anxiety through the secretion of ammonia bubbles by stressed individuals. Though the study of bird behaviour is at least partially knowable through visual means, the Five Domains model is not suitable for all animal taxa, like reptiles, for whom welfare indicators are difficult to gauge (Baker et al. 2013). Nor is it an option for understanding the materiality of plants or retrospectively, post-mortem, to infer how an organism lived. We know less about welfare conditions in personal ownership of birds, though Collard and Dempsey (2013) note that once in a new home, the animals’ lives are a shadow of their previous existence in the wild.

### Socioecological harm reduction interventions

(Termeer et al. 2019) highlight that only partial solutions are feasible for wicked problems, which the songbird trade certainly constitutes. Following (Hübschle and Margulies 2024), we propose a series of socioecological harm reduction approaches to songbird IWT that focus on reducing harm to birds without compromising livelihoods along the four stages of the supply chain (Figure 1A). In theory, these interventions construct a pathway to sustainable bird trade in the region. However, it is important to note these are only suggestions and ultimately the contours of any IWT intervention must be decided by local communities (Green 2025).

#### 1. Harvesting

‐ Assess species-specific mortality rates (e.g., corticosterone spikes (Cockrem 2007)) to list species too sensitive for harvesting.
‐ Implement payment schemes to release rarer species (trialed in shark and manta ray fisheries in Indonesia (Booth et al. 2023)).
‐ Phase out birdlimes due to damage to flight feathers (Platt et al. 2012).
‐ Promote peer-peer training from expert mist-net trappers and fund less lethal hunting methods like mist-nets, for poorer harvesters

#### 2. Transit

‐ Bocetti (1994) found that a quiet, dark environment with food, water, and minimal handling increases survival during transport for a Noth American insectivorous songbird. (Fiennes et al. forthcoming) note insectivorous birds dominate in Indonesia’s bird trade.
‐ Reduce overcrowding in transport vehicles (European Food Safety Authority 2006).
‐ Transport birds in their normal social groups (European Food Safety Authority 2006), where possible.

#### 3. Point of Sale

‐ Model conditions for expensive birds across all marketplaces: clean shops, spacious cages, food, and water.
‐ Install barriers between birds and visitors (Bartels et al. 2022).
‐ Reduce overcrowding in cages and avoid stacking birds (Bartels et al. 2022).

#### 4. Ownership

‐ Understand folk medicine practices by birdkeepers to promote avian longevity.
‐ Engage with aviculture communities and those interested in DIY aviary building to communicate welfare models like the five domains.

### Concluding remarks

To conclude, there are various structures of power (framed by economic and class issues) that impose violence on birds but also human actors to varying extents, such as those who rely on external investment or are tied to the patronage of more powerful actors. Here, we recognise and explore the importance of birds as mediums for socialisation and income and as status symbols for a range of men from different socioeconomic backgrounds in Indonesia.

By recognising songbirds as individual victims we have provided a more detailed understanding of the harms birds face in trade. Shifts towards the more sustainable use of birds (i.e. using less destructive harvesting and transport methods and at a rate that does not lead to long-term decline) can also reduce social harm to actors more vulnerable to economic inequalities (Lunstrum et al. 2023) in the songbird trade system.

Songbird trade is useful for case study of a more-than-human political ecology as it is embedded in complex and overlapping cultural systems (Javanese, footballing and more generic consumer cultures) and involves hundreds of species (Fiennes et al. 2024). But at the same time, the lessons we present here can be applied to other wildlife trades, including socially acceptable trades and those involving many species like invertebrates and plants. Ultimately, we hope this approach can be used to understand other illegal wildlife economies in terms of the social and cultural ideas that shape them, the material properties of traded entities, and how trade disrupts wild lives.

## Notes

### Competing Interest Statement

The authors have declared no competing interest.

### Funding

- Leverhulme Trust
- Society for Conservation Biology

