## Supporting Information for "A more-than-human political ecology of Indonesian songbird trade"

**Table 1** Comparison of the specifics of the Five Freedoms and the Five Domains Model. Reproduced from (Sopel 2021).

| The Five Freedoms | The Five Domains |
| --- | --- |
| 1. Freedom from hunger and thirst | 1) Nutrition- sufficient, balanced, varied and clean food and water |
| 2. Freedom from discomfort | 2) Environment – comfort through temperature, substrate, space, air, odour, noise and predictability |
| 3. Freedom from pain, injury and disease | 3) Health – enabling good health through the absence of disease, injury and impairment with a good fitness level |
| 4. Freedom to express normal behaviour | 4) Behaviour – providing varied, novel and engaging enrichment through sensory inputs, exploration, foraging, bonding, playing, retreating and others |
| 5. Freedom from fear and distress | 5) Mental state – the animal should benefit from predominantly positive states, e.g., pleasure or comfort, whilst reducing negative states such as fear, frustration, hunger, pain, or boredom |

**Table 2** Violations of the five domains towards birds in trade noted in the literature and hypothesized differences between the wild and captive lives of birds.

| Domain | Example |
| --- | --- |
| Nutrition | Some birds change colours in the marketplace, the Javan Green Magpie, *Cissa thalassina,* goes from green to blue from a lack of lutein (a yellow-colored pigment found in fruits, vegetables, and eggs) in its diet (Van Balen et al. 2013) |
|  | Lack of food and/or water in cages |
| Environment | As an example, the White-rumped Shama, *Copsychus malabaricus*, the territory is usually about 0,1 hectares (about 1.000 square metres). Area-wise, bamboo cages typically found in marketplaces are sometimes 30 x 30 cm; this is dramatically different from 1000 square metres. |
|  | Migratory birds fly hundreds of meters high, none of which will be accommodated even by the highest aviary in the vertical sense. |
|  | The harm birds may experience may also complicate their identification. For instance, their tails may abrade. Shorter tails are standard for birds in cages as the tail abrades really fast in the cage, and this is not a feature you can use in markets for identification (J. Eaton, pers. comms). |
|  | In German bird markets, Bartels et al. (2022) also recorded a lack of or insufficient protection from weather conditions like solar radiation (similar to Vignette 3). |
| Health | At Pramuka market, we saw broiler-style crates, which can cause osteoporosis in hens due to cramped conditions and lack of exercise, making their bones prone to breaking (Peng and Broom 2021). Nijman et al. (2022) found that these plastic crates were also used to transport Wandering Whistling-ducks, *Dendrocygna arcuata*, from Lake Jempang in Kalimantan, without room to move or access to food or water, for 15 to 20 hours by road, regularly violating basic welfare provisions. |
|  | In May 1994 at Pramuka bird market, a bird offered as a Critically Endangered Bali myna was actually a Black-winged Myna, *Acridotheres melanopterus* with its yellow eye-patches colored blue (Nijman et al. 2021) |
|  | Ferns (2017) also recorded dead and decomposing birds at the bottom of cages amongst the living birds in Sumatra marketplaces (Figure 1D) |
| Mental State | Birds, especially corvids, form strong social bonds and can experience grief. This suggests they might grieve the loss of their wild lives and social connections when in captivity (Dooren 2014). |

**Figure 1:** A) White-headed Munias, *Lonchura maja,* experiencing overcrowding. B) Evidence of mortality: a dead spotted dove, with two warblers in the same bin underneath other rubbish. C) Sunda Scops-owlets, *Otus lempiji* covered in or in close proximity to excrement. D) A dead munia in a water pot, E) Baby chickens spray painted. Photos taken by the first author.

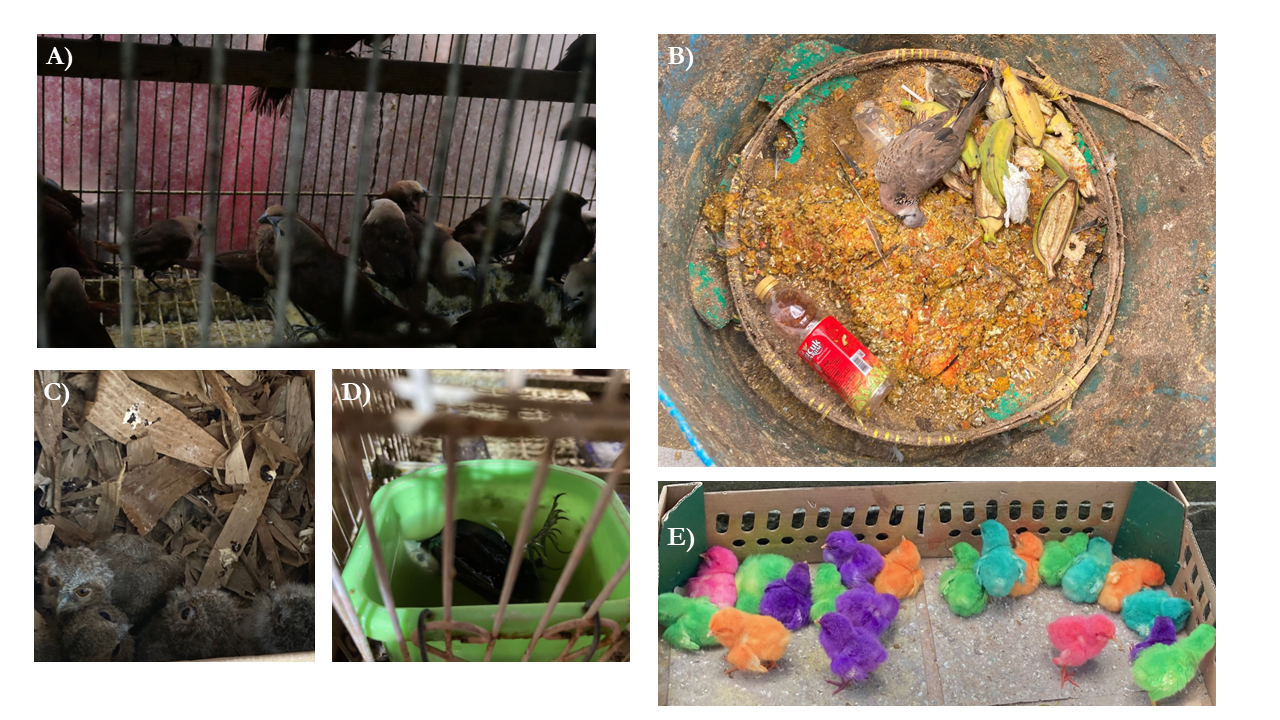

**Figure 2:** Blue-faced Honeyeater, *Entomyzon cyanotis* observed in a marketplace, potentially transported up to 5,000 km (photo by first author).
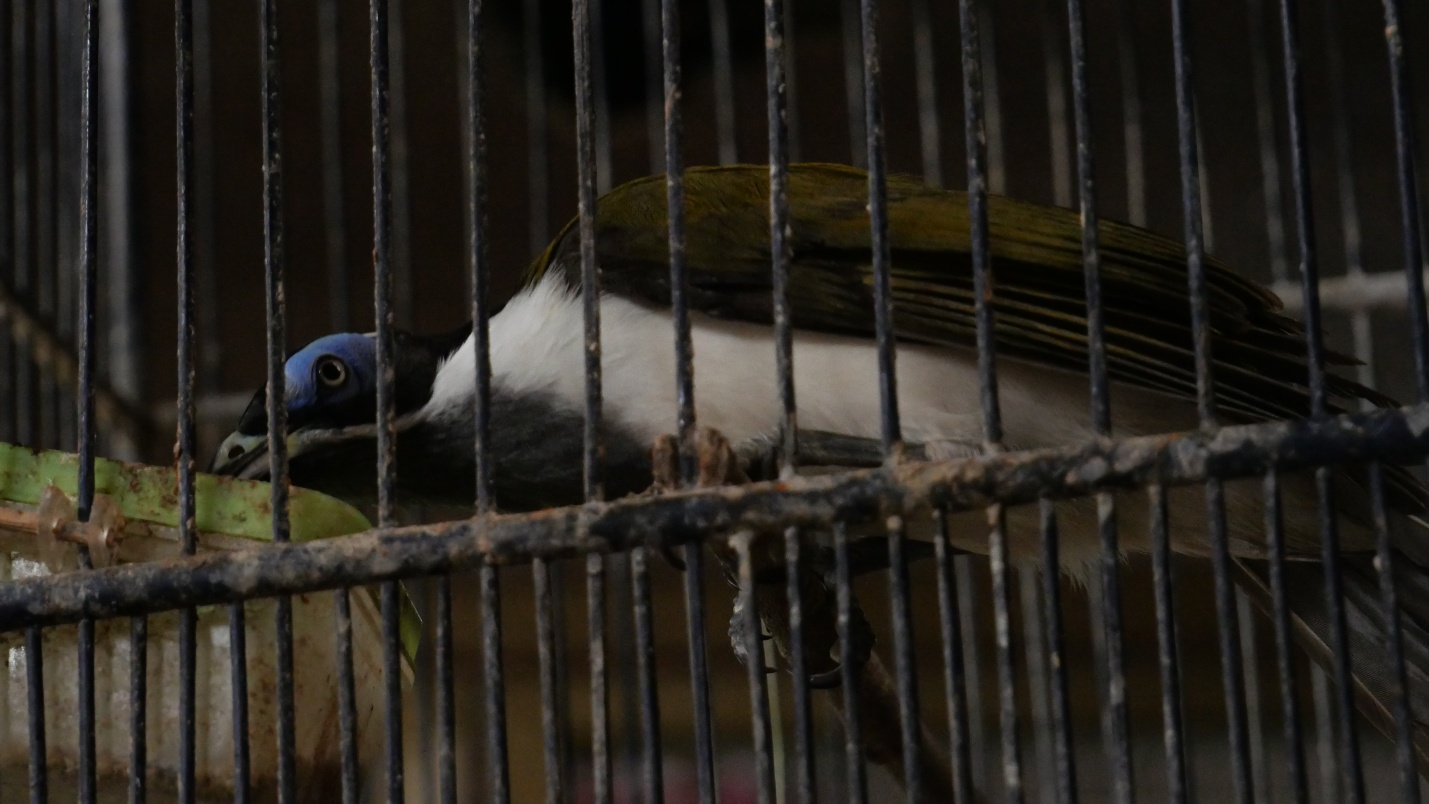

**Figure 3:** Other entities sold at wildlife marketplaces as multispecies communities; A) forest cats, b) civets and long-tailed macaques, C) live insects to feed insectivorous birds and D) flying squirrels. Photos taken by the first author.

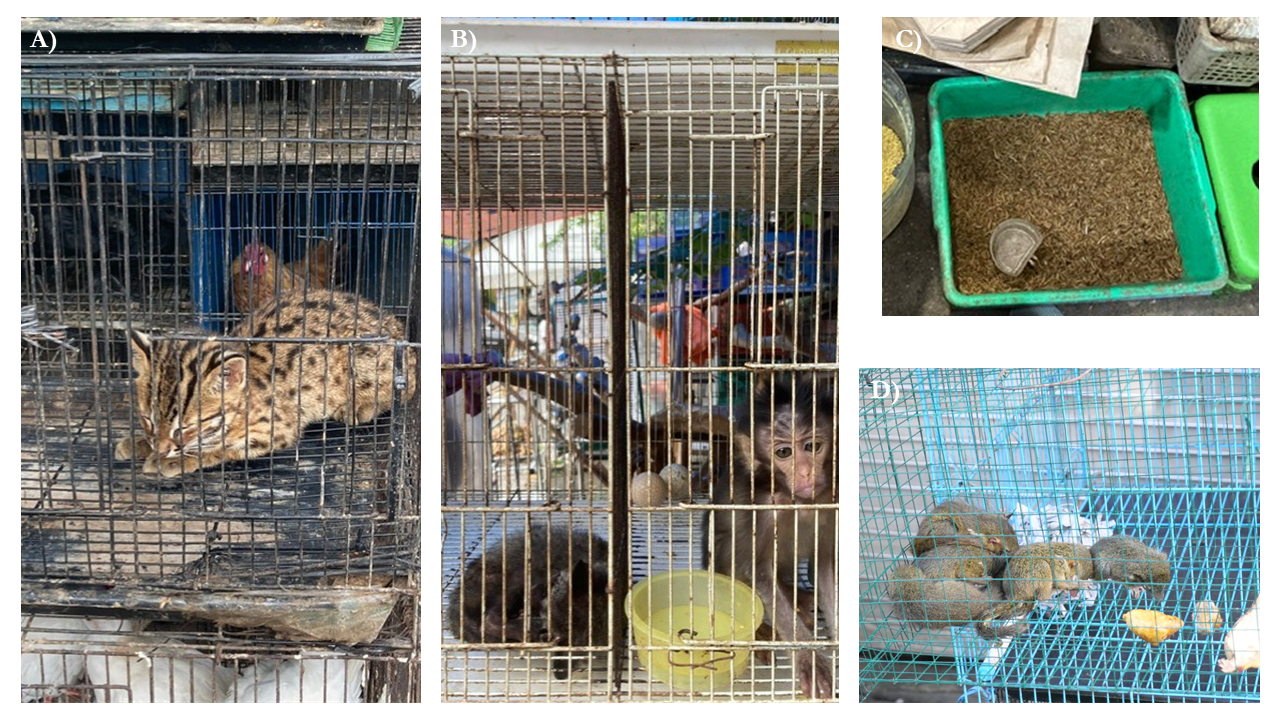

**Table 3**: Evidence from interviews (and the literature) for human trade actors.^[1]^

| **Stage** | **Role** | **Quotes** |
| --- | --- | --- |
| Harvester | Subsistence (Phelps et al. 2016) | Some people also hunt for themselves [for personal consumption] (ACL1) |
|  | Opportunist (Phelps et al. 2016) | Opportunistic hunting reflects societal norms where capturing and selling wildlife, whether from forests or urban areas, is normalized. People commonly take something from their backyard to sell or make money from it (ACL1) |
|  |  | In urban areas like Jakarta, people opportunistically catch birds, including escapees. They use traps with bait in high trees to capture species like barn owls, shamas, bulbuls, and shrikes, then sell them at markets or directly to buyers (ACL1) |
|  | Rule abuser (Phelps et al. 2016) | Lasia and other islands adopted the practice, leading to the probable extinction of the Nias Hill Myna. Palembang and Javan traders introduced it, and now locals are continuing it (ACI1) |
|  | Local guide (Phelps et al. 2016) | The locals had told us there were no more shamas left, as they had already caught them all. It seems the local island population learned these practices from Javan and city traders who visited and taught them (ACL1) |
|  | Specialist commercial (Phelps et al. 2016) | The islands of Lasia and Babi were overrun by about 1,000 traders, who gathered at the same coffee shop. This attracted more people, leading to widespread bird trading. As a result, islands like Lasia and Nias adopted this practice, causing the probable extinction of the Nias Hill Myna and the depletion of shamas. (ACI1) |
|  |  | With little left for dedicated trappers, small-scale efforts by local people are more common. They likely give birds to middlemen or local traders, such as bird shops in the capital town of Simeulue (ACI1) |
|  | Recreational (Phelps et al. 2016; Marshall et al. 2020) | Maybe not just small businesses, more like a side hustle. Sometimes they give them away, keep them for a while, and then sell them later (NGOL4) |
|  | Bycatch (Phelps et al. 2016) | In Java, particularly near markets like those in Bandung, small-scale hunters may catch birds in nearby forests, when they're hunting something else, and bring them to the market to sell (ACL1) |
| Intermediary | buying agents (Tsing 2015) | Middlemen go to the forest, pay local villagers to catch birds, and collect them through their own hunters. They receive orders from markets, consumers, or resellers, and then deliver the birds (ACL1, NGOL6) |
|  | Couriers (Guciano et al. 2024) | Sometimes transporters are given vehicles by investors. As a smuggler, I would drive the car from Sumatra to Java, but the car is owned by the investor. I receive bribe money for this role (NGOL8) |
|  |  | From Nusa Tenggara to East Java, birds are usually transported by trucks or ships with couriers, so they are not alone (GAKK-SUR-4) |
|  | Dropshippers (penampung)(Basuni and Setiyani 1989) | There’s another middleman who connects with buyers, including online shops and small-scale traders. The virtual trader can act as a physical marketplace, wholesaler, or sell specific types without a physical kiosk (ACL1, NGOL8) |
|  |  | Trucks transporting birds from Sumatra to Java also supply stalls in Pramuka, either as orders or stock. There are several middlemen involved, and the birds are distributed to small-scale traders, kiosks, Pramuka, and online sellers (ACL1) |
|  | specialist wholesaler | Trucks from Sumatra to Java follow a quota system and meet traders at locations like gas stations for bird drop-offs. These traders, referred to as middlemen, handle the distribution (ACL1) |
|  |  | Middlemen send birds to a penampung (collector/gatherer), where they are collected in a warehouse before being packaged and shipped to Java (ACL1) |
| Vendors | door-to-door sellers (Jepson et al. 2011) | Seen on motorbikes at Sukahaji (First author), also NGOL6 |
|  | wholesalers | As the owner of CV. Aneka Burung, I sold a cockatoo to a seller at the Pramuka bird market. Additionally, I have colibris (sunbirds), which are very cheap and can be sold directly to hawkers or consumers (NGOL6) |
|  | local shop owners (Jepson et al. 2011) | Seen in peri-urban areas in West Java and Kalimantan by the first author |
|  | bird shop (kisok) owners (Jepson et al. 2011) | NGOL6 |
| Consumers | Hobbyist (Marshall et al. 2020) |  |
|  | Contestant (Marshall et al. 2020) |  |
|  | Breeder (Marshall et al. 2020) |  |
|  | Affluent collector | Insight from authors |

**Table 4**: Methods for trapping and transporting birds from interviews and focus groups.

| **Stage** | **Method** | **Shortened** |
| --- | --- | --- |
| Trapping | bird lime | Opportunistic trappers use birdlime, typically derived from rubber or jackfruit sap (ACL3, NGOI2, NGOL4). If the bird is smooth and not stuck in the birdlime for long, the damage is minimal, and its feathers remain smooth (ACL3, NGOL7) |
|  |  | Birds caught using glue, adhesive, or birdlime often have damaged wings, they are not in high demand and don't fetch a good price (NGOL7) |
|  |  | I don't know why they use birdlime; I think it's an old traditional method (NGOI2) |
|  | mist nets | Mist nets are often used fishnets (NGOL7, ACL3) |
|  |  | back in the day, when Chestnut-capped Thrushes were trapped a lot in the lesser Sundas, the guys used to stick up mist nets about 200 metres long. Mistnets are definitely the most common (NGOI2) |
|  |  | they'll [harvesters] just stick 20 metre poles either side [of the net] and then just stick it up… they put it in the breakaways between the trees, in clearings, over roads (NGOI2) |
|  |  | Mist nets are quite expensive, costing around USD 50 (NGOI2) |
|  | canopy mist nets | Trappers use canopy nets to catch entire flocks, which has increased the trade of canopy species like minivets and nuthatches (NGOI2) |
|  | fish hook | hooks, fishhooks (NGOL4) |
|  | bird to scare off | But I think people are more often using owls to attract. Yeah, and other birds to sort of, to then scare the owl off. You might get a mixed flock coming in to sort of sort of do it or whatever (NGOI1) |
|  | poison | termites on the hook (NGOI1) |
|  | baited cages | They set up cages in the canopy with fruit to trap Hill mynas, using a rope to hoist the open cage. I'm unsure how effective this method is (ACI1) |
|  |  | Hunters typically use traps with trails that birds follow, leading them to get trapped (BKSDA_4) |
|  | playback lures | That's created a massive problem online, Sound Recording libraries… are now restricted, but it didn't always used to be like that' ACA_OUT_MAL_1 |
|  | decoy birds | A singing male bird attracts another for a contest in the forest. Hunters use a two-cage system, placing a male from their collection on the ground to attract another male or a mate. This creates a natural competition, aiming to capture a wild male. The cage is modified to close when the bird enters to eat |
| Transport | woven palm leaf pouch | Birds can be brought to hunters in leaf packages, specifically inside ketupat (woven palm leaf pouches). These are made when the bird comes from the hunter and is then taken to the middleman in the forest (NGOL5) |
|  | socks | When my friends patrolled the forest, hunters used "kaos kaki" (socks) or fabric bags with strings to transport birds. They tied the beak with tape, wrapped the bird in fabric or newspaper to protect its wings, and fixed the legs to pre0vent breakage. They meticulously placed the birds in socks, tying and hanging them. One, two, three, four, five hanging there… (NGOL7) |
|  | beaks tied with tape | When my friends patrolled the forest, hunters used "kaos kaki" (socks) or fabric bags with strings to transport birds. They tied the beak with tape, wrapped the bird in fabric or newspaper to protect its wings, and fixed the legs to prevent breakage. They meticulously placed the birds in socks, tying and hanging them. One, two, three, four, five hanging there… (NGOL7) |
|  | cloth bags | The birds in the bag, like the shama bird or Oriental magpie-robin, are expensive and classy. They are rarely put in socks and are usually placed in small cages (5x5 or 10x10). The bags are rotated and modified. Hunters still use this method, packing the birds tightly so they remain silent (NGOL7) |
|  |  | I think the cloth bag is most common when you go into the forest anyway. Yeah, you just sort of tied them off to your bag or whatever and dangle them (NGOI1) |
|  | In shoeboxes | They used to transport the birds in shoeboxes, poking holes for air (NGOI2) |
|  | watermelons | The bird might be repackaged or disguised in different ways. When it goes to the middleman and then to consumers, the packaging varies. Some use watermelons. Some use wire, sticks, and skewers to prevent sagging, with 5 to 10 birds inside. However, the birds become wet immediately (NGOL7) |
|  | wooden cages | for expensive ones (ACL1) |
|  |  | The birds in the bag, like the shama bird or Oriental magpie-robin, are expensive and classy. They are usually placed in small cages (5x5 or 10x10) (NGOL7) |
|  | thermoses | One bird per thermos (NGOL7) |
|  | plastic poultry crates | Observed by authors in marketplaces |
|  | cardboard boxes | I saw two young traders posing with a ruffled female leafbird (possibly a blue-winged, Javan, or greater-green leafbird) that came from a cardboard box (Observed by authors in marketplaces) |
| Sale | cement bags | Observed by authors in marketplaces |
|  | cages | Observed by authors in marketplaces |
|  | plastic bags | Observed by authors in marketplaces |

**Figure 4:** Birds traded in football-branded cages; A) a canary, *Serinus canaria* in an FC Barcelona cage and B) a Crimson Sunbird, *Aethopyga siparaja* in an Arsenal cage.

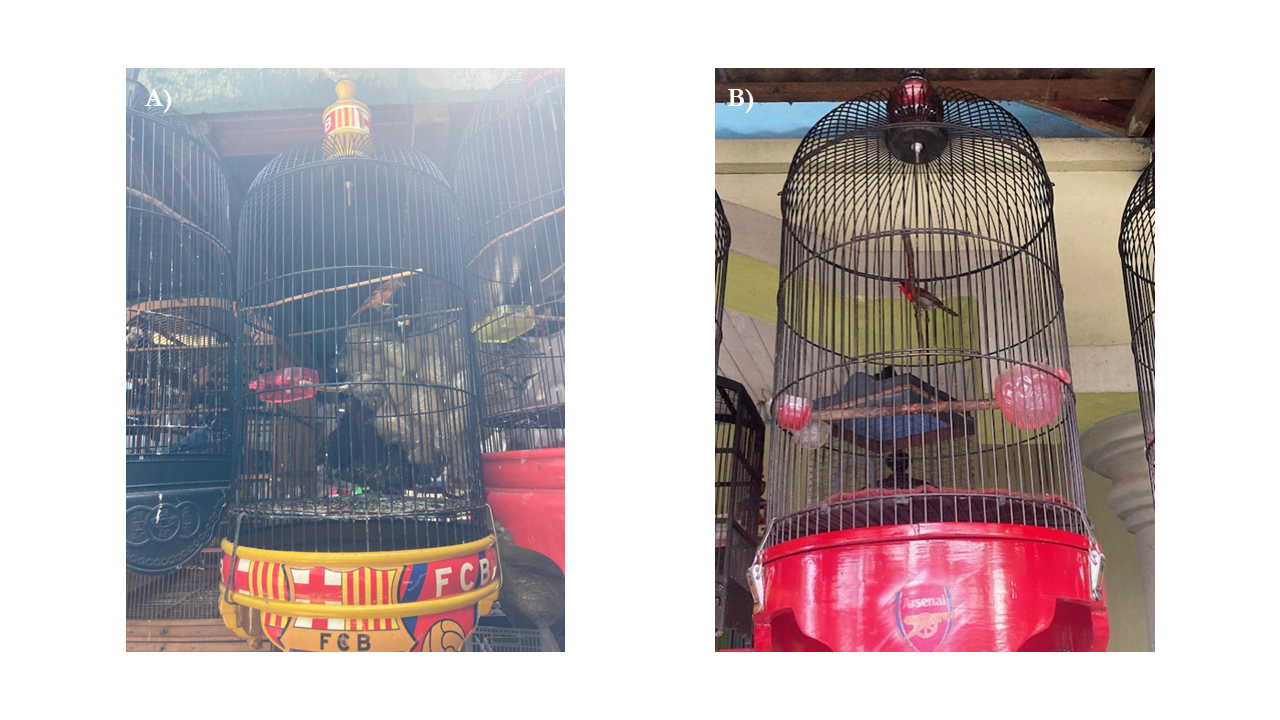

**Guide 1: Focus group structure and script** **for workshops with Indonesian conservation law enforcement, in English and Indonesian.**

When referring to participants, the codes for conservation law enforcement are BKSDA, GAKK, KAR, and POL, followed by a number (e.g., "BKSDA_2"); "X" is used for unidentified participants. Academic (ACA) and NGO (NGO) codes are location-based (e.g., "ACAI1" for an international academic and "ACAL2" for an academic local to Indonesia).

The workshops were structured in five stages:

1. Short introduction and project presentation – *empathy and understanding*
2. Icebreaker exercise using audiovisual montage - *sensory elicitation*
3. Asking law enforcement about their job in general and different enforcement activities that they are engaged in – *focus group*
4. The wider context of their job – the environment in which they operate and how they relate to birds – *focus group*

The questions in these stages were structured as follows:

**2.  Icebreaker exercise- watching the audiovisual materials of the marketplace to elicit a sense of closeness to the marketplace and setting the scene**

Participants were asked the following questions:

- What did you think of that? Any feelings or responses you would like to share please write on the board/flipchart
- What is your experience watching the audiovisual materials?

**3.  Asking law enforcement about their job in general and different enforcement activities that they are engaged in (what happens when they see a protected species, what is the process of species identification that they currently use)**

- What three things are helpful for your job, and what three things could hypothetically make your job easier? Please write these thoughts on a sticky note and put them up on the board
- How would you feel if you were to discover a market trader who is selling endangered/protected species?
- Follow-up: how would you feel to discover that this bird is now endangered in the wild?

**4. The wider context to their job – the environment in which they operate and how they relate to birds**

- What do birds mean to you?
- What do you think are the potential consequences of certain bird species going extinct?
- Do you think extinction is an important issue?
- Follow-up: How do you feel about this?
- Follow-up: Are you aware of any traded bird species that are threatened?
- Do you think popular songbird species are declining in Indonesia*?* Follow-up: What are the reasons for the decline?

**5. Narratives about technology in their work/CLE attitudes to technology and the use of technology in their job**

- How do you currently identify birds you encounter in your work and how effective are these methods?
- If they do not currently use any technology: Why are you currently not engaging with any digital tools to do this?
- If they are using technology already for songbird identification: What does this technology do well and what does not work? If you could redesign existing technology, what techniques would you use?

**Guide 2:** Interview structure and script (a selection of questions relevant to this paper, part of a broader interview on experiences visiting wildlife marketplaces and bird trade) for discussions with academics and conservation organization staff for the purposes of triangulating the focus group data.

**PERSON EXPERIENCE VISITING WILDLIFE MARKETPLACES**

1. How often do you go to markets?
2. Since what year, when was the first time you ever went to one?
3. How many wildlife markets have you ever been to in your life?
4. And if species do go extinct from trade, what do you think is the impact on bird trade or bird keeping practices?
5. in terms of sustainability and welfare aspects of trade, what's your experience of this and your opinion on the state of trade relating to mortality and welfare issues?
6. How do you feel about keeping birds in cages?
7. What is the future of bird trade in Indonesia?
8. What do you personally feel and what do you think is, is realistic or achievable?

**TRADE CHAIN AND THE STATUS IN GENERAL**

**Enforcement within the trade chain**

9. Do you think the current methods of law enforcement for songbird’s work? So, this could be like the law itself, or activities related to the law (or more generally about the effectiveness of the law, for example)
10. How do you think bird trade should be enforced, in an ideal world?

**IDENTIFICATION**

11. How would you describe your identification capabilities?
12. What methods if any, do you use for species identification?
13. What helped you get better at identification?

1. Role references are included in the column . in the role section, if a name comes from existing literature, it is referenced. Otherwise, terms were used by interviewees or conservation law enforcement officers.
